# *Listeria monocytogenes* cell-to-cell spread bypasses nutrient limitation for replicating intracellular bacteria

**DOI:** 10.1101/2025.01.31.635960

**Authors:** Prathima Radhakrishnan, Julie A. Theriot

## Abstract

*Listeria monocytogenes* is an intracellular bacterial pathogen that obtains nutrients from the mammalian host cell to fuel its replication in cytosol. Sparse infection of epithelial monolayers by *L. monocytogenes* results in the formation of distinct infectious foci, where each focus originates from the initial infection of a single host cell followed by multiple rounds of active bacterial cell-to-cell spread into neighboring host cells in the monolayer. We used time-lapse microscopy to measure changes in bacterial growth rate in individual foci over time and found that intracellular bacteria initially replicate exponentially, but then bacterial growth rate slows later in infection, particularly in the center of the infectious focus. We found that the intracellular replication rate of *L. monocytogenes* is measurably decreased by limiting host cell glucose availability, by decreasing the rate of intracellular bacterial oligopeptide import, and, most interestingly, by alterations in host cell junctional proteins that limit bacterial spread into neighboring cells without directly affecting bacterial growth or metabolism. By measuring the carrying capacity of individual host cells, we found that the nutritional density of cytoplasm is comparable to rich medium. Taken together, our results indicate that the rate of intracellular *L. monocytogenes* replication is governed by a balance of the rate of nutrient depletion by the bacteria, the rate of nutrient replenishment by the metabolically active host cells, and the rate of bacterial cell-to-cell spread which enables the bacteria to seek out “greener pastures” before nutrient availability in a single host cell becomes limiting.

**SIGNIFICANCE:** *Listeria monocytogenes* is a foodborne pathogen that is capable of facultative intracellular growth in a wide variety of mammalian cell types. It uses actin-based motility to spread directly from the cytoplasm of one infected host cell into another. After initially infecting cells in the small intestine, *L. monocytogenes* can circulate inside of infected macrophages, and subsequently spread to cells of distal organs, where it can cause spontaneous abortions in pregnant women and meningitis in immunocompromised individuals. In addition to facilitating body-wide dissemination, the actin-based cell-to-cell spread of *L. monocytogenes* contributes to the ability of the pathogen to evade the cell-mediated arm of the immune system. In this work, we find that cell-to-cell spread also benefits the bacteria in a third way, by enabling the bacteria to overcome the nutrient limitation they would face if they remained confined to a single host cell, such that they never enter stationary phase.

## INTRODUCTION

Growth of bacteria in culture is well-defined and can be well-characterized by simple rules (Monod, 1949; Neidhardt, 1999). When inoculated into fresh media, most bacteria exhibit a lag phase where the growth rate is close to zero. As bacteria adapt to their environment, they enter an acceleration phase, where growth rate increases until it reaches the exponential phase, when bacteria are replicating at a constant rate. Next, bacterial growth rate begins to decline due to the depletion of nutrients in the media or the build-up of toxic metabolites, followed by a stationary phase where the net growth rate returns to zero as the rate of bacterial replication is balanced by the rate of cell death. Finally, cultures enter a death phase where the absolute numbers of bacteria decline as the rate of cell death exceeds the rate of replication (Monod, 1949).

Under ideal conditions when nutrients are not limiting, bacterial growth can be modeled surprisingly well using the simple exponential function (Neidhardt et al., 1999):

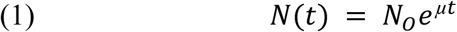

which can be rewritten as:

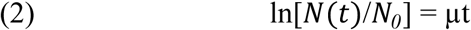

where *N(t)* represents the bacterial number at time *t*, *N_0_* the initial number of bacteria in the culture, and *µ* the constant growth rate expressed in units of inverse time. For a wide variety of bacterial species under a plethora of experimental conditions, it has been observed that the growth rate, *µ*, correlates with various features of the bacterial cell, including cell size, chemical composition, and rates of transcription and translation (Schaechter et al., 1958; Neidhardt, 1990; Neidhardt, 1999; Scott et al., 2010; Erickson et al, 2017; Si et al., 2019; Belliveau et al., 2021). The consistent relationship between the growth rate for bacterial cells and their physical characteristics is remarkable, and corroborates the view that bacterial growth follows simple, quantitative rules (Neidhardt, 1999).

The more complex reality where the rate of bacterial growth changes over time as a function of culture stage is typically modeled phenomenologically using sigmoidal functions that require the introduction of additional quantitative parameters, though none of these functions includes the final death phase (Pearl and Reed Lowell, 1920; Zwietering et al., 1990; Wachenheim et al., 2003). One such function that has been widely used to model changes in bacterial growth rate as a function of nutrient availability (Atolia et al., 2020) is the Gompertz model (Zwietering et al., 1990; Tjorve et al., 2017):

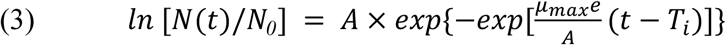

where *N(t)* is the number of bacteria as a function of time, *N_0_* is the initial number of bacteria in the culture, *µ_max_* is the maximal growth rate (units of inverse time) observed during exponential phase, *A* is the “carrying capacity” or maximum number of bacteria that can be supported at stationary phase, and *T_i_* is the time of the inflection point where the growth rate begins to slow down. At very long times, this equation asymptotes to the carrying capacity A, and exhibits its maximum growth rate *µ_max_* at the time of the inflection point T_i_. Although the Gompertz model can be expressed in a variety of mathematical forms (Gompertz et al., 1825; Laird et al., 1964; Gibson et al., 1987; Zwietering et al., 1990; Tjorve et al., 2017), this particular form has the advantage that the constants *A*, *µ_max_* and *T_i_* can all be directly measured from experimental data.

*Listeria monocytogenes* is a facultative intracellular bacterial pathogen (Gray and Killinger, 1966; Vázquez-Boland et al., 2001; Gray et al., 2006). After invading a mammalian host cell, the pathogen escapes from the phagosomal compartment (Portnoy et al., 1988; Marquis et al., 1995), replicates within the cytoplasm of the host cell (Freitag et al., 2009; Chen et al., 2017) and upregulates the transcription of virulence genes (Mengaud et al., 1991; Chakraborty et al., 1992; Freitag et al., 1993). In the host cell cytosol, the bacterium uses actin-based motility to move through the cytoplasm of the host cell (Dabiri et al., 1990; Theriot et al., 1992) and spread from one cell to the other without escaping into the extracellular space (Tilney and Portnoy, 1989; Robbins et al., 1999). Outside of infected hosts, *L. monocytogenes* can be easily cultured in liquid media under laboratory conditions (Jones and D’Orazio, 2013). It is also frequently found growing in ready-to-consume foods (Gray et al., 2004), and in soil or silage where it feeds off of decaying plant material (Welshimer, 1960; Freitag et al., 2009; Lang Halter et al., 2013). While growth of *L. monocytogenes* in culture is known to progress along the six phases outlined by Monod (Bruno and Freitag, 2011; Jones and D’Orazio, 2013), the dynamics of bacterial growth in the cytoplasm of infected host cells have not been studied in comparable detail.

A growth curve of *L. monocytogenes* replicating inside of mammalian host cells might or might not resemble that of bacteria growing in culture. While *L. monocytogenes* utilizes host cell nutrients for intracellular growth (Chen et al., 2017), bacterial replication within an intracellular niche has not been demonstrated to deplete the host cell of nutrients in the way that bacterial replication is known to deplete liquid media of nutrients over time. In particular, the host cell metabolism can continue to process and synthesize nutrients throughout the course of infection, and *L. monocytogenes* has even been shown to increase the host cell’s glycolytic activity (Gillmaier et al., 2012). Moreover, intracellular *L. monocytogenes* can escape their immediate environment altogether and access fresh nutrients by spreading into neighboring cells.

We sought to measure intracellular growth of *L. monocytogenes* in mammalian host cell epithelial monolayers over time using live cell imaging. Time-lapse imaging of live cells expressing a fluorescent protein allows the direct observation of growth in individual bacterial foci that originate from a single infected host cell over long time periods, without reducing the viability of either the bacteria or the host cells (Robbins et al., 1999; Ortega et al., 2019; Bastounis et al., 2020). This method also enables us to monitor changes in growth rates over time upon perturbation of virulence behaviors such as cell-to-cell spread. Overall, our results demonstrate that the net rate of *L. monocytogenes* replication in the cytosol of mammalian host cells depends on the competition between the rate of nutrient depletion by growing bacteria with the rate of nutrient replenishment based on host cell metabolic activity and the rate of accessing new nutrient pools via bacterial spread into naive, uninfected host cells. By continually spreading into new host cells, rapidly growing *L. monocytogenes* are able to maintain their density below 2% of the maximum host cell carrying capacity.

## RESULTS

### Intracellular Growth Rate of *Listeria monocytogenes* Decreases Over Time

To measure changes in the rate of intracellular bacterial growth over time, we collected long-term time-lapse movies of *L. monocytogenes* strain 10403S expressing red fluorescent protein (RFP) propagating in a monolayer of A431D human epidermoid epithelial cells. Host cells were infected using a low density of bacteria, such that only about 1 in 10^3^ cells in the monolayer were invaded, resulting in well-spaced individual foci that could be followed over time. Bacterial foci were imaged at five-minute intervals over ten hours, beginning at 5 hours post-infection (h.p.i) and continuing until 15 h.p.i., using a wide-field microscope objective with a low numerical aperture (20X, N.A. 0.75) so that individual bacteria at any vertical position within the host cells relative to the coverslip would be captured within the focal plane in a single image (Fig. 1A). Illumination conditions and imaging frequency were optimized so that essentially no photobleaching occurred over the time scale of observation. From 5 to 15 h.p.i., foci expanded in all directions, with irregular shapes that varied among foci (Fig. 1B-D). In order to quantify the number of bacteria in each image, we used total RFP fluorescence intensity as a proxy for bacterial number. Early in each time-lapse sequence, individual bacterial cells were easily identified as discrete fluorescent points, and we confirmed that integrated fluorescence intensity was strongly correlated with bacterial number (Supp. Fig. 1A). Although individual bacteria were no longer distinguishable at late time points, total bacterial fluorescence intensity continued to increase smoothly and monotonically.

**Figure 1:**
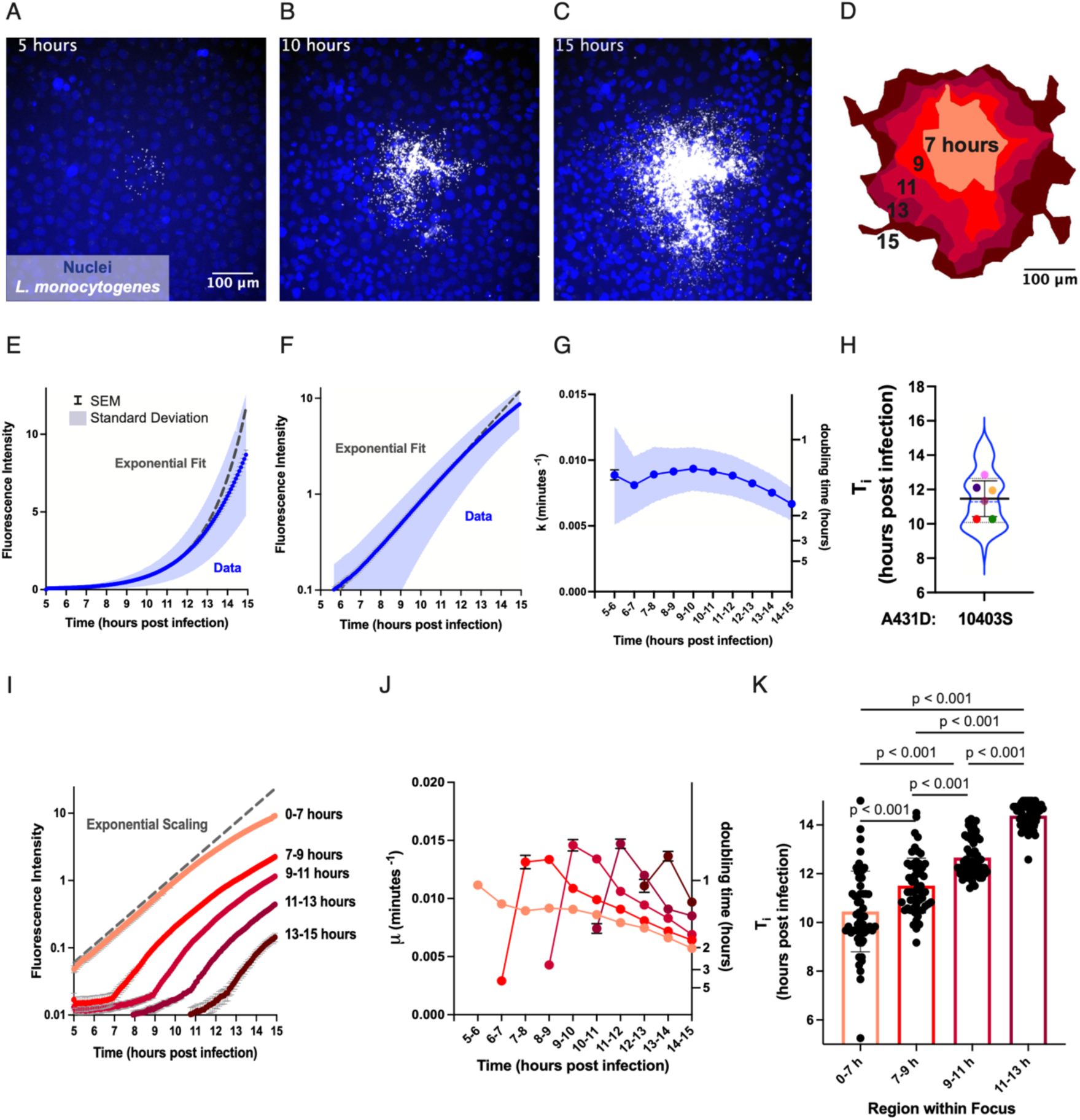
Intracellular growth rate of *L. monocytogenes* decreases over time dependent on distance from the origin of the focus. A-C. Epifluorescence microscopy images of nuclei (Hoescht, blue) and intracellular *L. monocytogenes* (mTagRFP, gray) at 5 (A), 10 (B) and 15 (C) hours post infection. D. Schematic representation of bacterial focus shown in A-C. The outer limit of each region represents the maximum intensity projection of the bacterial focus from the beginning of the time-lapse movie to the labeled time. (See Supp. Fig. 2A and Materials and Methods for details) From the center out, the regions include all positions occupied by bacteria from 0-7 h.p.i., 7-9 h.p.i., 9-11 h.p.i., 11-13 h.p.i. and 13-15 h.p.i. E. Quantification of average bacterial fluorescence intensity as a function of time for 140 foci. The dashed line shows the fit to the portion of the growth curve depicting exponential growth at a constant rate. F. Quantification of average bacterial fluorescence intensity as a function of time for 140 foci on a semilog plot. As in the previous panel, dashed line depicts the fit to the portion of the curve describing exponential behavior. G. Growth rate calculated within 1 hour time intervals as a function of time for 125 foci. For panels E-G: the error bars represent the standard error of the mean and the shaded region the standard deviation. H. Inflection point (T_i_) at which bacterial growth begins to slow for 122 foci recorded over six independent experiments. Each colored dot represents the average of one independent experiment. Thick black lines show the mean, whiskers show the standard deviation, dashed line represents the median, and dotted lines the first and third quartiles. I. Fluorescence intensity as a function of time for each region shown in panel D. Data from 58 foci was averaged for each temporally defined region. Error bars represent S.E.M. J. Growth rate as a function of time for bacteria enclosed in each region for 67 foci. K. Inflection point (T_i_) at which bacterial growth begins to slow for bacteria in each region of the focus as shown in D. Each point represents a single focus. Error bars depict standard deviation and p values were calculated using the Wilcoxon Rank-Sum test. For panels I-K: The color of each trace or bar corresponds to a region on the schematic shown in D.

We portrayed change in bacterial growth rate over time by constructing a growth curve, where total bacterial fluorescence intensity was plotted against time (Fig. 1E-F). Intracellular bacterial growth was initially exponential (Neidhardt et al., 1999) as described by Equation (1), over the time span from 5 h.p.i. to about 11 h.p.i. (Fig. 1E). When the vertical axis was converted to a log scale, the portion of the growth curve representing constant exponential growth appeared to fit a straight line as expected (Fig. 1F). During this early stage of growth, the average growth rate 𝜇 across 122 foci as defined by Equation (2) was 0.0087 min^-1^ (S.D. = 0.0026 min^-1^), equivalent to a doubling time of about 80 minutes. However, the growth curve for all observed *L. monocytogenes* foci began to noticeably deviate below the exponential fit after about 11 h.p.i. (Fig. 1E-F).

Next, we calculated the value of the apparent growth rate constant over time by fitting distinct exponential functions to subsets of the growth curve at one-hour time intervals (Supp. Fig. 1B-C, see *Methods*). On average, the apparent growth rate stayed fairly constant at around 0.009 min^-1^ until about 10-13 hours post-infection, and then began to decline (Fig. 1G). This behavior is reminiscent of bacterial growth in batch culture, where exponential growth with a constant growth rate is followed by a slow-down of growth rate as the increasing number of bacteria depletes the media of nutrients and causes the accumulation of toxic metabolites (Monod 1942; Monod, 1949). To determine when intracellular bacterial growth rate begins to fall, we determined the inflection point *T_i_* as defined in Equation (3) for each individual focus. To this end, we fit distinct exponential functions to increasing numbers of fluorescence intensity measurements, starting at the first frame of the time-lapse recording and adding one data point from the following frame with every iteration until we fit the entire growth curve to an exponential function in the last iteration (Supp. Fig. 1D). We then calculated the growth rate constant 𝜇 for each fit and defined the time at which 𝜇 began to decrease monotonically, which we identified as the inflection point *T_i_*. The average inflection point for the 122 foci was 11.5 hr (S.D. = 1.6 hr) (Fig. 1H).

### Time of Bacterial Growth Slow-down Increases with Distance from the Origin of the Focus

Because each bacterial focus starts from a single infected host cell and then propagates outward through the epithelial monolayer, host cells near the center of the focus are expected to become densely crowded with *L. monocytogenes* earlier than those at the periphery. To determine whether a decline in bacterial growth rate follows the attainment of high bacterial cell density within individual host cells, we sought to monitor change in bacterial growth rate over time in various positions within a single focus.

To this end, we partitioned each focus into five regions, and determined the growth rate µ and inflection point T_i_ as described above, for each region. The first region included all positions occupied by *L. monocytogenes* from the initial time of infection to 7 h.p.i. The second region covered the additional area occupied by bacteria from 7 to 9 h.p.i., the third from 9 to 11 h.p.i., the fourth from 11 to 13 h.p.i. and the last from 13 to 15 h.p.i. (Fig. 1D). These regions were generated by drawing tight outlines called alpha shapes (Edelsbrunner et al., 1983) around maximum intensity projections of the bacterial focus from the start of the time-lapse recording to the appropriate time points set two hours apart (Supp. Fig. 2A). On average, bacteria grew exponentially in the most interior region with an initial value of 𝜇 = 0.009 min^-1^, until the growth rate began to decline at an average inflection point at 9.8 h.p.i. (S.D. = 1.6 h.p.i.) (lightest shade in Fig. 1 I-K).

When we monitored growth rate in the outer regions, we consistently found the apparent initial value of 𝜇 to be significantly higher than the expected 0.009 min^-1^ (Fig. 1J). Using a computational stochastic simulation of *L. monocytogenes* cell-to-cell spread (Ortega et al., 2019), we found that these inflated initial apparent values of 𝜇 could be attributed to active actin-based movement of *L. monocytogenes* from the dense center of the focus outward toward the periphery (Supp. Fig. 2B-C). After accounting for this phenomenon, we were able to accurately determine an inflection point T_i_, where the observed growth rate began to decline, for every region of each focus (Fig. 1K; see *Methods*). The inflection point time increased significantly with every consecutive region moving outward from the center of the focus, such that the growth rate began to decline earliest in the most interior region and later in each subsequent region (Fig. 1K). This observed delay in inflection point timing as a function of distance from the center of the focus is consistent with our hypothesis that increasing bacterial cell density over the course of infection causes a slowing down of bacterial growth rate, such that the growth rate shows first in the central (oldest) regions of the focus.

### Nutrient Deprivation Restricts Intracellular Growth of *L. monocytogenes*

*L. monocytogenes* utilizes peptides and carbohydrates present in the host cell cytosol as energy sources for intracellular growth (Borezee et al., 2000; Chico-Calero et al., 2002; Joseph et al., 2008; Eylert et al., 2008; Grubmüller et al., 2014). In particular, pathogenic *L. monocytogenes* imports sugar phosphates that are products of glucose metabolism, such as glucose-6-phosphate, to fuel cytosolic growth (Chico-Calero et al., 2002). Furthermore, glucose availability has been found to limit growth of *L. monocytogenes* in milk and in liquid media (Pine et al., 1989). To test the hypothesis that nutrient restriction might contribute to the observed decrease in intracellular bacterial growth rate over time, we exchanged full media for glucose-free media at 5 h.p.i., the time at which we began collecting time-lapse movies of bacterial growth (Fig. 2A). Bacterial growth rate from 5 to 6 h.p.i. was similar between the two conditions, indicating that glucose availability within the host cell cytoplasm was sufficient to support robust bacterial growth within this time interval. However, the value of µ declined rapidly beginning at 6 h.p.i. in glucose-free media but (as described above) remained fairly constant in full media until approximately 11 h.p.i (Fig. 2B). The magnitude of the decrease in growth rate was also more severe in glucose-free as compared to full media (Fig. 2B) and the number of bacteria per focus was much lower in the glucose-free condition as compared to full media by the final time point at 15 h.p.i. (Fig. 2A). These results led us to conclude that nutrient deprivation in the host cell environment can cause a decrease in growth rate of *L. monocytogenes* over time.

**Figure 2:**
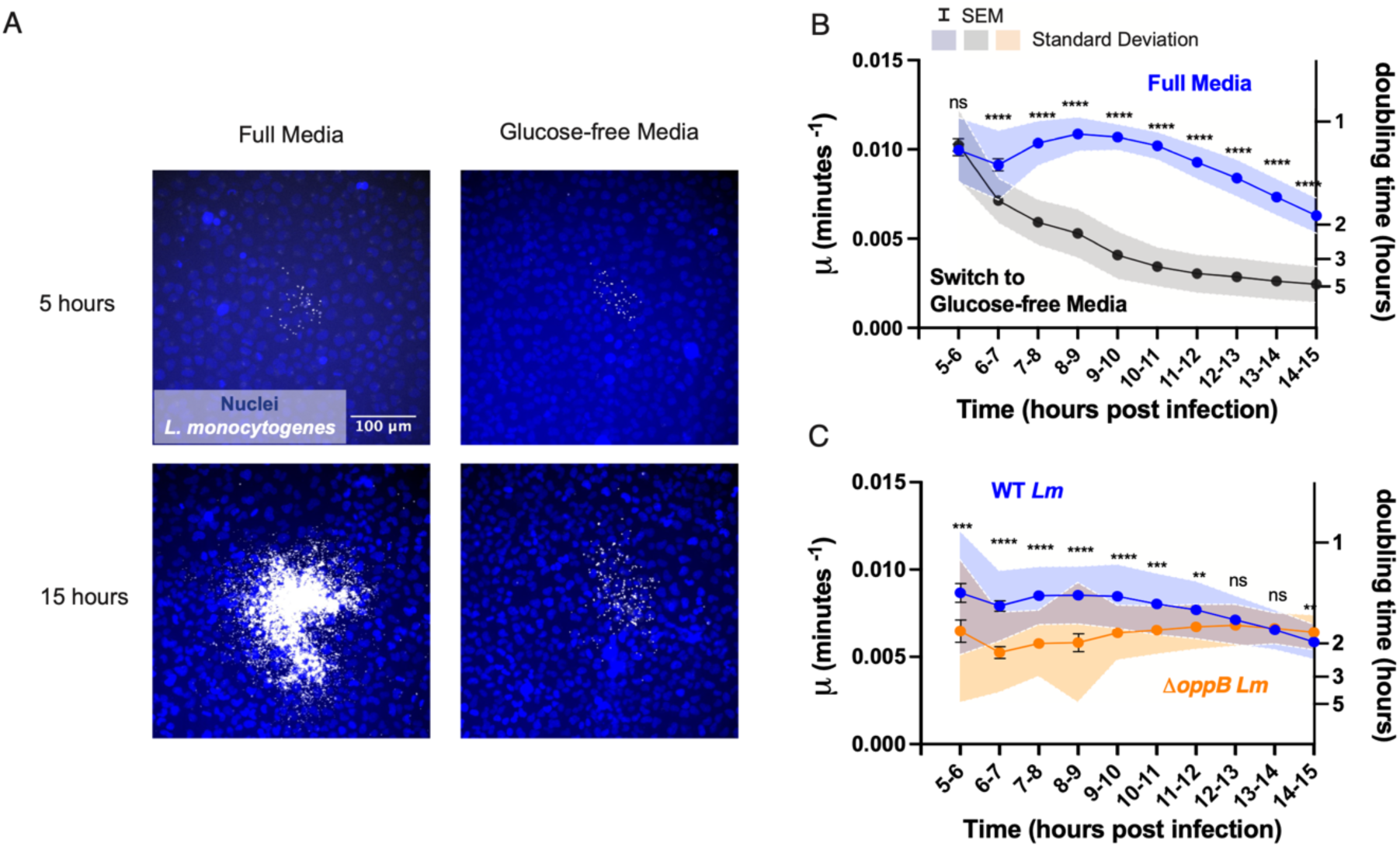
Nutrient deprivation restricts intracellular growth of *Listeria monocytogenes*. A. Micrographs depicting nuclei (Hoescht, blue) and intracellular bacteria (mTagRFP, gray) at 5 (top row) and 15 (bottom row) h.p.i. Cells were immersed in full media in the left column for the full length of the experiment. Cells in the right column grew in full media until 5 h.p.i. when full media was exchanged for no glucose media. B. Growth rate as a function of time for cells immersed in full media (blue trace) and cells growing in no glucose media beginning at 5 h.p.i. (black trace). 34 and 33 foci were imaged for each condition respectively. C. Growth rate as a function of time for WT *Listeria monocytogenes* compared with growth rate as a function of time for *ΔoppB Listeria monocytogenes*. 43 and 44 foci were imaged for each condition respectively. Error bars represent the standard error of the mean and shaded area the standard deviation. Significance of the difference between grown rates of each condition (full media vs. no glucose media in B. and WT *Listeria monocytogenes* vs. *ΔoppB Listeria monocytogenes* in C.) were tested at one hour time intervals. P-values were calculated using the Wilcoxon Rank-Sum test and denoted as asterisks above each data point (see Methods. “n.s.” = not significant).

### Intracellular Growth Rate of a Slowly-Replicating *L. monocytogenes* Mutant Does Not Decrease Over Fifteen Hours of Infection

*L. monocytogenes* is an amino acid auxotroph; that is, it cannot synthesize all 20 amino acids used for protein production, and typically acquires amino acids from the host cell cytoplasm (Siddiqi and Khan, 1989; Premaratne et al., 1991; Eylert et al., 2008). Oligopeptides containing between 2 and 10 amino acid residues serve as the main amino acid source for intracellular growth of *L. monocytogenes* (Marquis et al., 1993; Verheul et al., 1995; Verheul et al., 1998; Borezee et al., 2000). To internalize oligopeptides from the host cell, the bacterium employs the oligopeptide permease Opp system, which includes OppB, an integral membrane protein component of the transporter (Borezee et al., 2000). Due to its inability to efficiently access nutrients required for growth, we would expect Δ*oppB L. monocytogenes* to exhibit a slower intracellular growth rate as compared to the WT bacterium (Borezee et al., 2000). Indeed, we found that, between 5 and 6 h.p.i., growth rate of Δ*oppB L. monocytogenes* replicating within A431D cells was significantly reduced in comparison to that of WT *L. monocytogenes*, averaging around 0.006 rather than 0.009 min^-1^ (Fig. 2C).

When we monitored change in bacterial growth rate over time, we found that, while growth rate of WT *L. monocytogenes* began to decline after 11 h.p.i., Δ*oppB L. monocytogenes* grew at a fairly constant (slow) rate up until at least 15 h.p.i., the full period of our observations (Fig. 2C). As there are fewer Δ*oppB* bacteria in any infected cell as compared to WT *L. monocytogenes* at a given moment, this result is consistent with our hypothesis that slowing of bacterial growth occurs when the number of cytoplasmic bacteria reaches a threshold at which nutrient availability limits further growth. In this scenario, because the slow-growing Δ*oppB* bacteria never pass this threshold, their growth rate is never forced to decrease. In other words, the observed slowing of the intracellular replication rate for *L. monocytogenes* around 11 h.p.i. is most likely to be caused by a competition between the rate of nutrient utilization by the bacteria and the rate of nutrient replenishment by the metabolically active host cells.

In addition to nutrient limitation, quorum sensing has been shown to reduce the metabolic activity of other intracellular pathogens like *Burkholdaria glumiae* at high bacterial cell densities by restricting glucose uptake (An et al., 2014). However, we did not observe a difference in either the inflection time *T_i_* or the net change in growth rate µ between WT *L. monocytogenes* EGD-e and a quorum sensing mutant, Δ*agrD L. monocytogenes* (Supp. Fig. 3). Therefore, quorum sensing by the *agr* system is unlikely to contribute to any decrease in growth rate that might result from accumulation of *L. monocytogenes* within infected host cells.

### Restricting *L. monocytogenes* Cell-to-Cell Spread Slows Bacterial Growth at Late Time Points in Infection

Next, we sought a way to increase the number of cytoplasmic bacteria within individual host cells without altering the bacterium’s physiology or level of metabolic activity. To do so, we restricted *L. monocytogenes* cell-to-cell spread through various perturbations, such as expressing E-cadherin mutants in host A431D cells that lack the cytoplasmic domain (Δcyto E-cadherin) or contain two point mutations that prevent them from being ubiquitinated by the ubiquitin ligase, hakai (K738R, K816R E-cad) (Radhakrishnan et al., 2022). Due to the increased difficulty of escaping into a neighboring host cell, we would expect intracellular bacteria to reach the threshold density at which growth slows at an earlier time when cell-to-cell spread is limited as compared to control conditions. To determine how reducing bacterial spread by altering host cell-cell junctions alters bacterial growth rate over time, we compared time-lapse movies of *L. monocytogenes* spreading through a monolayer of WT or K738R, K816R E- cad A431D cells (Fig. 3A-B). As previously reported (Radhakrishnan et al., 2022), this host cell mutation results in significantly smaller infectious foci as compared to *L. monocytogenes* infection of WT host cells (Fig. 3C). Although this very subtle host cell perturbation is not expected to have any direct effect on bacterial growth or metabolism, the final number of bacteria in infectious foci at 15 h.p.i. was also substantially reduced as compared to foci in A431D cells expressing normal E-cadherin (Fig. 3D).

**Figure 3:**
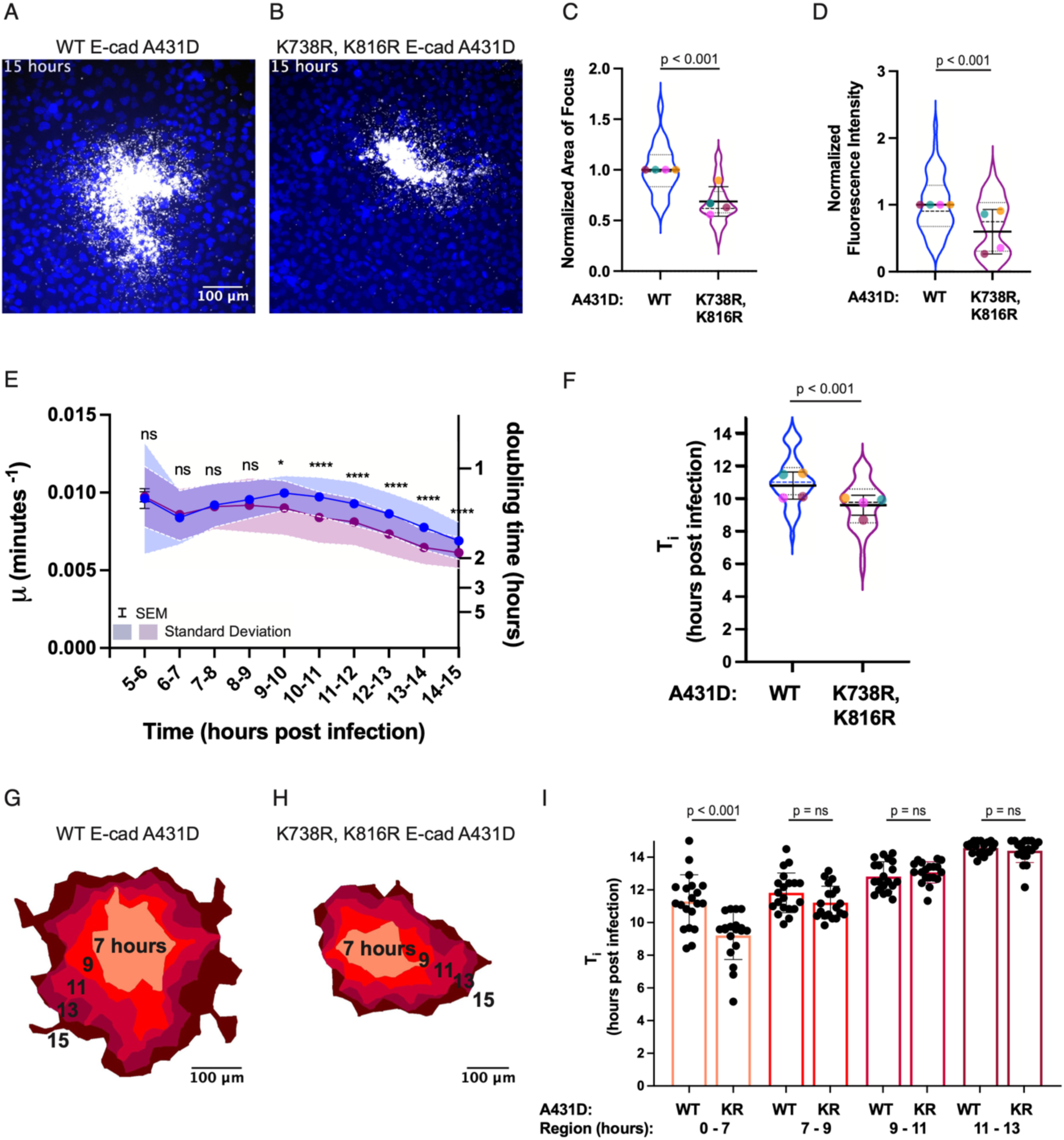
Intracellular growth rate of *L. monocytogenes* begins to slow earlier when actin-based cell-to-cell spread is decreased. A-B. Micrographs depicting *L. monocytogenes* (mTagRFP, gray) propagating in a monolayer of WT E-cad (A) or K738R, K816R E-cad (B) A431D cells. C-D. Comparison of focus area (C) and fluorescence intensity per focus (D) at 15 h.p.i. between WT E-cad (blue violin) and K738R, K816R E-cad (purple violin) A431D background. Each colored dot represents the average of one independent experiment. Thick black lines show the mean, whiskers show the standard deviation, dashed line represents the median, and dotted lines the first and third quartiles. E. Growth rate as a function of time for bacteria propagating through WT E-cad (blue trace) and K738R, K816R E-cad (purple trace) A431D cells. F. Comparison of inflection points (T_i_) for bacteria replicating in WT E-cad or K738R, K816R E-cad cells. C-F. 50 and 45 foci were imaged for each condition respectively. P values for C, D and F were determined using the linear mixed-effects model and p values for E were calculated using the Wilcoxon Rank-Sum test and denoted as asterisks above each data point (see Methods; “n.s.” = not significant). G-H. Schematic representations of bacterial foci depicted in A. and B. with each focus partitioned into 5 regions as in Figure 1D. I. Inflection point (T_i_) at which bacterial growth begins to slow in each region of the focus in WT or K738R, K816R E-cad A431D cells. 20 foci for each condition were included in this analysis. The color of each bar corresponds to the region on the schematics in E and F. P values were calculated using the Wilcoxon rank-sum test.

The value of 𝜇 at the beginning of the recording (5 h.p.i.) was comparable between the two conditions, suggesting that this perturbation to cell-to-cell spread does not alter the initial rate of bacterial replication during the exponential phase. However, bacteria in K738R, K816R E-cad cells deviated from exponential growth earlier than *L. monocytogenes* spreading through WT E-cad cells, resulting in an earlier inflection time T_i_ (Fig. 3E-F). Bacterial growth rate µ also fell to a lower value in the K738R, K816R E-cad background as compared to the WT E-cad background during the time of observation (Fig. 3E). These results are consistent with our hypothesis that increased bacterial cell density due to the hindrance of cell-to-cell spread causes a greater decrease in bacterial growth rate over time as compared to conditions where spread is unrestricted.

Next, we partitioned each bacterial focus into five regions as described previously to enable direct comparison of bacterial growth as a function of location (Fig. 3G-H). The inflection point T_i_ in the most interior region, which includes all positions *L. monocytogenes* occupied from 0-7 hours of infection, was significantly earlier in the K738R, K816R E-cad background as compared to the WT E-cad background (Fig. 3I). However, there was no significant difference between bacterial growth rate in WT and K738R, K816R E-cadherin backgrounds for every subsequent region, which represent positions visited by the bacteria from 7 to 15 hours post infection (Fig. 3I). As an independent confirmation of these results, we performed similar measurements for *L. monocytogenes* growing in host cells with the Δcyto E- cad background and found essentially identical results (Supp. Fig. 4). .

Overall, if bacterial growth rate slows upon achievement of a threshold density of bacteria or degree of nutrient consumption per host cell, we would expect to observe fewer bacteria per focus in conditions where cell-to-cell spread is limited and a larger number of bacteria per focus in conditions where cell-to-cell spread is enhanced. In addition to the E-cadherin mutations detailed above, several other host cell perturbations are known to decrease *L. monocytogenes* cell-to-cell spread and thereby decrease focus size, including inhibition of caveolin-mediated endocytosis and inhibition of macropinocytosis (Radhakrishnan et al., 2022). Conversely, treatment of host cells with the growth factor EGF is observed to enhance *L. monocytogenes* cell-to-cell spread and increase focus size (Radhakrishnan et al., 2022).

When we re-analyzed data that was previously reported in Radhakrishnan et al., 2022 by measuring the fluorescence intensity per focus, we discovered a lower bacterial number in comparison to the control at 12 h.p.i. in multiple conditions where *L. monocytogenes* cell-to-cell spread is impaired, and an increase in bacterial number per focus upon EGF treatment (Fig. 4A-C). Across all conditions tested, the bacterial number appeared to correlate strongly with focus size (Fig. 4C). Because these treatments all target the host cell to influence the efficiency of bacterial cell-to-cell spread, rather than targeting bacterial metabolism or replication directly, we conclude that the simplest explanation for this strong correlation is that *L. monocytogenes* replication in a single infected host cell is necessarily limited until the bacteria can access fresh sources of nutrients by spreading into uninfected neighboring cells.

**Figure 4:**
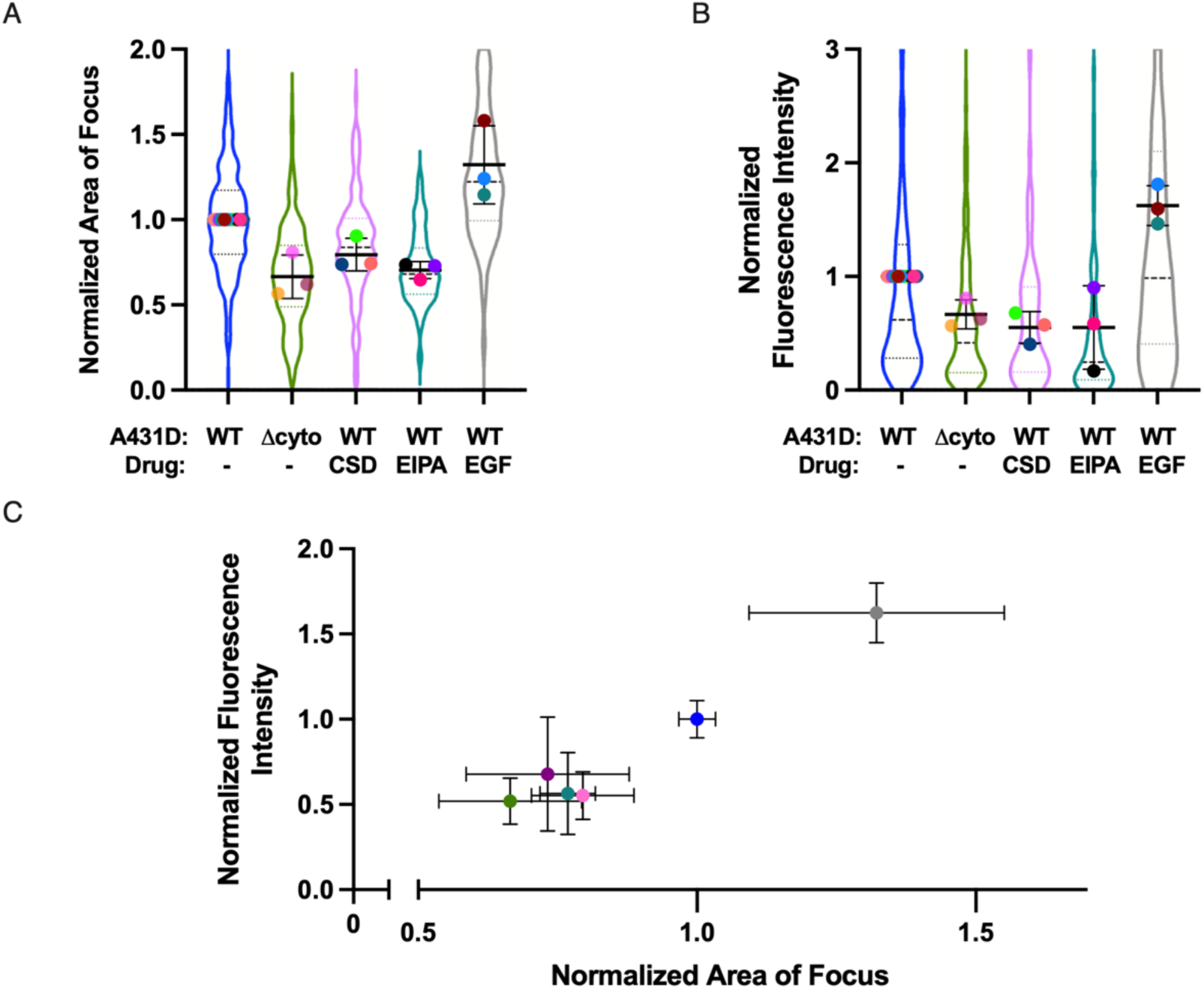
Focus size and total bacterial number decrease or increase when the efficiency of cell-to-cell spread is restricted or enhanced. A-B. Comparison of focus area (A) and fluorescence intensity per focus (B) at 12 h.p.i. between control conditions (blue violin), Δcyto E-cad A431D background (green violin), CSD treatment in WT E-cad A431D cells (pink violin), EIPA treatment in WT E-cad A431D cells (teal violin) and EGF treatment in WT E-cad A431D cells (gray violin). 448, 235, 140, 141 and 143 foci were imaged for each condition respectively. Each colored dot represents the average from one independent experiment. Thick black lines show the mean, whiskers show the standard deviation, dashed lines show the medians, and dotted lines the first and third quartiles. The p-value was determined using the linear mixed-effects model (see Materials and Methods). C. Scatter plot depicting focus area values in A. plotted against fluorescence intensity values in B. The color of the data points in C corresponds to the color of the violins in A and B and represents the same condition. Whiskers show standard deviation.

### The Carrying Capacity of Mammalian Host Cell Cytoplasm Is Comparable to Rich Medium

In order to measure the maximum density of bacteria at stationary phase in an infected mammalian cell (that is, the carrying capacity, indicated as *A* in Equation (3)), we used a bacterial mutant incapable of performing actin-based cell-to-cell spread. In this strain, designated as “mutant 34” in the initial report, five adjacent basic residues (KKRRK) in the ActA protein-coding sequence have been replaced with alanine, rendering the mutant strain incapable of nucleating actin filament growth (Lauer et al., 2001). Because this mutant strain expresses a full-length version of the ActA protein under its normal genetic regulation, the overall metabolic requirements of this strain are expected to be nearly identical to wild-type bacteria. In order to measure the volume of a single host cell accurately during the period of bacterial replication, we transfected A431D cells with a plasmid expressing the green fluorescent protein mEmerald, at a low multiplicity, such that about 1% of cells expressed soluble mEmerald, filling the cytoplasm. In order to increase the range of cell sizes in the epithelial monolayer, we co-expressed the VSV-G glycoprotein, which mediates cell-cell fusion (Florkiewicz and Rose, 1984, Feliciano et al., 2018). In cultured monolayers, we could readily identify individual unfused mEmerald-expressing cells as well as much larger fused cells forming syncytia with multiple nuclei, ranging in area up to more than ten-fold the size of unfused cells (Fig. 5A-B). Using confocal microscopy, we found that the height of A431D cells in these confluent monolayers was quite constant at 11 µm ± 2 µm (mean ± SD) for both fused and unfused cells (Fig. 5C), so we were able to accurately approximate the volume of every mEmerald-expressing host cell in every video frame over the time course of observation by measuring the projected area in each frame. Typically, individual A431D cells grew by about 35% (± 21%) during the 10 hours of observation.

**Figure 5:**
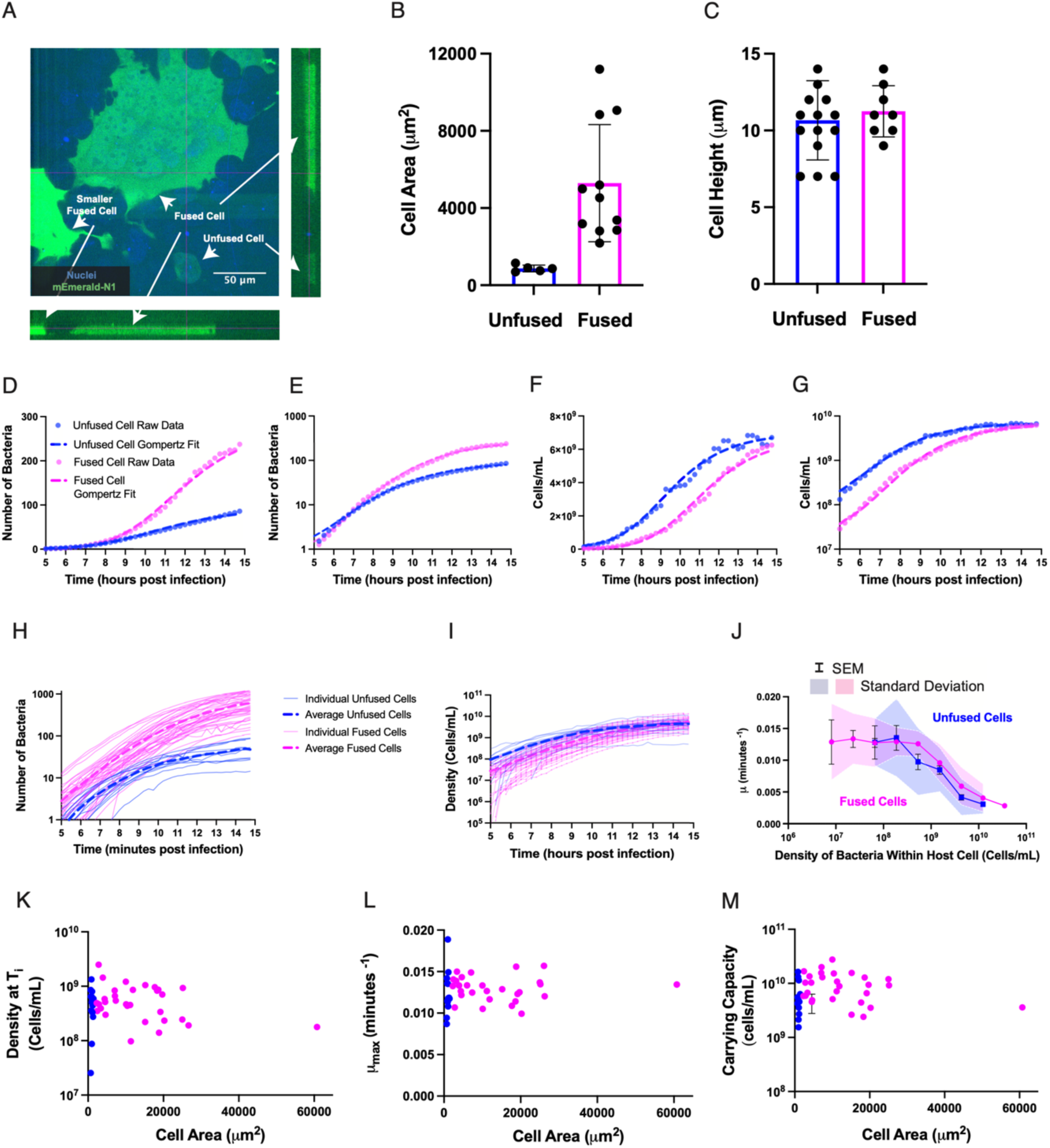
Intracellular bacterial growth rate decreases with increasing bacterial density, independent of host cell size. A. Orthogonal views of fused and unfused, transfected A431D cells with labeled nuclei (Hoescht, blue) and expression of a fluorescent cloning vector mEmerald-N1 (mEmerald, green) in transfected cells. Standard (x-y) view is shown at upper left, x-z cross-section at lower left, and y-z cross section at upper right. Cross-sections were sampled at locations indicated by thin magenta lines in the x-y view. Several mEmerald-expressing cells are labeled to indicate their positions in the orthogonal views. B. Range of total cell areas for unfused, transfected cells (blue bar) and fused, transfected cells (magenta bar). C. Cell heights (approximated to the nearest micron) for transfected, unfused cells (blue bar) and transfected, fused cells (magenta bar). For panels B and C: Error bars represent the standard deviation. P values for all panels were calculated using the Wilcoxon rank-sum test. D-E. Quantification of total bacterial number as a function of time for one representative unfused host cell (blue dots) and one representative fused host cell (magenta dots) shown on a linear plot (D) and semilog plot (E). Dashed lines show the fit of the growth curves to the Gompertz function. F-G. Quantification of intracellular bacterial density as a function of time for the same unfused host cell (blue dots) and fused host cell (magenta dots) as shown in D and E, represented on a linear plot (F) and semilog plot (G). Dashed lines show the fit of the growth curves to the Gompertz function. H-I. Bacterial number (H) or bacterial density (I) as a function of time for 14 infected unfused cells (blue traces) and 39 infected fused cells (magenta traces). Light, solid lines represent individual cells, while thick, dashed lines represent the average of unfused (blue line) or fused (magenta line) cells. J. Bacterial growth rate as a function of density within a host cell, where bacterial densities have been grouped into 10 bins that are equally spaced in the log scale. Growth rate values have been sorted into one of 10 bacterial cell density bins. The number of cells analyzed per condition include 14 unfused cells (blue trace) and 39 fused cells (magenta trace). Error bars represent standard error of the mean and shaded regions represent the standard deviation. K-M. Bacterial density at inflection point (T_i_) when bacterial growth begins to slow (K), maximal growth rate (L) and carrying capacity estimated by fitting growth curves to the Gompertz function (M) for 14 unfused host cells (blue dots) and 31 fused host cells (magenta dots). The error bars in panel M depict the standard error of the fits.

We infected these monolayers with the mutant 34 strain of *L. monocytogenes* and searched for individual cells (or fused syncytial cells) expressing mEmerald that also carried internalized *L. monocytogenes*. As expected, the bacteria grew to fill the cytoplasm over the period of observation (5-15 h.p.i.) without spreading into adjacent cells. We measured bacterial growth using total fluorescence intensity as described above. Growth curves for individual infected cells, both fused and unfused, exhibited easily observable deviations from exponential growth, appearing to slow down as they approached stationary phase. Notably, the total number of bacteria at the end of the observation period was substantially higher in the larger fused cells than in the smaller unfused cells (as shown for representative examples in Fig. 5D-E). However, when we converted bacterial number into bacterial density (cells/mL) by dividing the number by the total cell volume at each time point, we found that the final bacterial density in stationary phase was quite similar for both fused and unfused cells (Fig. 5F-G). Across a population of 14 unfused cells and 39 fused cells, we found consistent results where the absolute *number* of bacteria per cell at the end of the period of observation covered a wide range of values, but the final bacterial *density* was tightly distributed (Fig. 5H-I).

From the individual growth traces, we were able to estimate the growth rate *µ* as a function of bacterial density for all individual infected host cells, both unfused and fused. For both categories, the growth rate was fairly constant at low densities and began decreasing monotonically at densities above about 10^9^ cells/mL (Fig. 5J). The individual growth curves were well-fit by the Gompertz model (Equation (3)) (Fig. 5D-G). Using this fit, we were able to estimate the quantitative parameters of the model as a function of bacterial density for each trace, including the maximum growth rate (*µ_max_*), the density at inflection time (*T_i_*), and the density at full carrying capacity (*A*). Over the full range of observations including 14 individual unfused cells and 29 fused cells, we found that these parameters were essentially constant as a function of cell volume, even though the largest fused cells in this data set were about 60-fold larger than typical unfused cells (Fig. 5K-M). These observations strongly suggest that bacterial density, rather than absolute number of bacteria or time since infection, is the major determinant of bacterial growth behavior inside the cytoplasm of infected host cells.

Averaged over all infected host cells (both fused and unfused), the carrying capacity we observed for *L. monocytogenes* in mammalian host cell cytoplasm was 8.8 x 10^9^ cells/mL (SD = 5.7 x 10^9^ cells/mL). We note that some fraction of the host cell volume may not be accessible for bacterial growth (for example, *L. monocytogenes* do not typically grow inside of the nucleus), so this value may slightly underestimate the true carrying capacity of mammalian cytosol. Notably, this value is very similar to the maximal density reported for *L. monocytogenes* grown in rich medium (Brain Heart Infusion (BHI) broth) in stationary phase, about 5 x 10^9^ cells/mL (Bruno and Freitag, 2011).

Importantly, the highest density reached by wild-type bacteria in the center of the normal infectious foci over the time course of our observations, up to 15 h.p.i., never reaches, or even approaches, this carrying capacity limit. Over the 122 wild-type foci we observed, the maximal bacterial density in the central region was equivalent to 2.1 x 10^8^ cells/mL (SD = 0.7 x 10^8^ cells/mL), about 50-fold lower than the cytoplasmic carrying capacity. That is, wild-type bacteria can avoid having their growth restricted by nutrient limitation simply by spreading into neighboring cells.

## DISCUSSION

In his 1949 review titled, “The Growth of Bacterial Cultures”, Jacques Monod wrote, “The growth of bacterial cultures, despite the immense complexity of the phenomena to which it testifies, generally obeys relatively simple laws, which make it possible to define certain quantitative characteristics of the growth cycle… the accuracy, the ease, the reproducibility of bacterial growth constant determinations is remarkable and probably unparalleled, so far as biological quantitative characteristics are concerned.” (Monod, 1949; Neidhardt, 1999).

In this work, we have shown that, in spite of the additional complexities of growing within the cytoplasm of mammalian host cells and undergoing an elaborate process of actin-based motility to drive cell-to-cell spread, the intracellular growth of *L. monocytogenes* recapitulates key features of their growth in culture, and can be represented by highly reproducible quantitative parameters. At early times after infection, when nutrients are abundantly available, bacteria grow exponentially within host cell cytoplasm. As infection progresses and the metabolic activity of bacterial cells alters the composition of host cell cytoplasm, the bacterial growth rate slows. However, unlike growth within a culture tube, intracellular bacteria are able to escape into a neighboring cell and access fresh nutrient sources. They can then grow exponentially once more within the new host cell and begin the cycle anew.

In support of this model, we have shown that bacterial growth rate begins to slow earliest at the center of the infectious focus, where host cells are more densely populated with bacteria as compared to cells at the periphery of the focus. By partitioning a bacterial focus into distinct regions based on time of exposure to bacterial cells, we additionally demonstrate that the initial growth rate upon entry into uninfected host cells via cell-to-cell spread in the epithelial monolayer is similar to the rate of bacterial growth early in infection. Furthermore, bacterial growth in each subsequent region is exponential until a later time at which growth rate declines. The time of this inflection point time (T_i_), at which bacterial growth starts to slow down, occurs later and later with distance from the origin of the focus.

Our observation that bacterial growth rate slows earlier in glucose-free as compared to full media indicates that nutrient deprivation can reduce intracellular bacterial growth rate.

Moreover, we have shown that a slowly replicating *L. monocytogenes* mutant (Δ*oppB*) remains in exponential phase for a longer time as compared to the wild-type control. As bacteria are expected to consume nutrients at a rate which is proportional to their growth rate, this result is also consistent with a role for nutrient deprivation in the transition from exponential to stationary phase. However, we do not rule out other factors that might also play a role, such as greater accumulation of toxic metabolites (Nguyen and Portnoy, 2020) in the host cell Rcytoplasm with an increase in bacterial load. In any case, the spread of intracellular bacteria into neighboring host cells apparently serves to keep their density well below the nutrient-limited carrying capacity in that environment, by about a factor of 50.

The ability of *L. monocytogenes* to spread to neighboring cells is a trait that has been honed by evolution and is regulated by virulence genes (Lobel et al., 2012; Vasanthakrishnan et al., 2015; Haber et al., 2017; Krypotou et al., 2019). Our results showing a correlation between bacterial growth rate and efficiency of cell-to-cell spread suggest that intracellular bacterial growth involves a balance between the rate of nutrient depletion by the replicating bacteria, the rate of nutrient replenishment by the metabolically active host cells and the rate of bacterial spread to uninfected host cells. Under normal circumstances for wild-type bacteria, these rates appear to be closely matched, such that small decreases in bacterial growth rate (such as for the Δ*oppB* mutant) are sufficient to prevent any observable slowdown of bacterial growth over our 15-hour observation period, and conversely small decreases in the efficiency of bacterial cell-to-cell spread (as in the K738R, K816R E-cad A431D host cells) tip the balance such that the growth rate of intracellular bacteria slows down slightly earlier than for *L. monocytogenes* in wild-type host cells. As PrfA activation is required for efficient cell-to-cell spread (Shetron-Rama et al., 2002), our results demonstrate that, in addition to enabling dissemination through a monolayer without exposure to the host’s humoral immune response (Zenewicz and Shen, 2007), occupying a virulent state increases fitness of *L. monocytogenes* within its intracellular niche by matching the time-scale of spread to “greener pastures” to the time-scale associated with nutrient depletion inside of individual host cells.

## ACKNOWLEDGMENTS

We gratefully acknowledge Fabian Ortega for initial observations of *L. monocytogenes* growth slow-down in the center of infectious foci, and for numerous insightful scientific discussions, as well as for providing the simulation code used in Supplemental Fig. 2B-C. We thank Daniel Portnoy for providing the Δ*oppB* strain of *L. monocytogenes*, and Christian Riedel for providing the Δ*agrD* strain. This work was supported by NIH R37 AI036929 and by the Howard Hughes Medical Institute. This article is subject to HHMI’s Open Access to Publications policy. HHMI laboratory heads have previously granted a nonexclusive CC BY 4.0 license to the public and a sublicensable license to HHMI in their research articles. Pursuant to those licenses, the author-accepted manuscript of this article can be made freely available under a CC BY 4.0 license immediately upon publication.

## MATERIALS AND METHODS

### Bacterial Strains and Growth Conditions

The WT 10403S strain of *Listeria monocytogenes* was used for the majority of experiments in this study. To prevent visualization of extracellular bacteria, *L. monocytogenes* expressed mTagRFP under control of the *actA* promoter, which is only expressed once the bacterium enters the host cell cytosol (Zeldovich et al., 2011). This strain was created by transforming the plasmid pMP74RFP into *Escherichia coli* SM10 λpir. Then, the plasmid was delivered to *L. monocytogenes* by conjugation and integrated into the tRNA^ARG^ locus of the bacterial chromosome (Lauer et al., 2002, Ortega et al., 2017). The *actA* mutant *L. monocytogenes* that was used to infect transfected A431D cells was created by introducing the plasmid pMP74RFP into chromosomal *actA* mutant 34 (Lauer et al., 2001), via conjugation.

Specifically, amino acids 146-KKRKK-150 of ActA were changed to 146-AAAAA-150 in the creation of this mutant (Lauer et al., 2001). *ΔoppB L. monocytogenes*, a gift from Daniel Portnoy, included an in-frame deletion of *oppB* (Whiteley et al., 2015) in mTagRFP-expressing WT *L. monocytogenes*.

*L. monocytogenes* EGD-e::pAD_1_-cGFP, a gift from Christian Riedel, was created as previously described (Balestrino et al., 2010). Briefly, the *L. monocytogenes* integrative pH-*hly*- *gfp*-PL3 plasmid, which is under control of the Hyper SPO1 constitutive promoter (p*hyper*) was engineered to allow for certain elements to be easily replaced. Then, the *gfp* gene in pH-*hly*-*gfp*- PL was exchanged for the gene encoding GFP mut2 (Cormac et al., 1996), which exhibits higher fluorescence intensity. After the gene sequence was confirmed to be correct, the new plasmid, pAD_1_-cGFP, was transformed into *L. monocytogenes* EGD-e using electroporation.

Chromosomal deletion of *agrD* was conducted as described previously (Riedel et al., 2009, Monk Ian et al., 2008) to create *L. monocytogenes* EGD-e *ΔagrD*::pAD_1_-cGFP.

To prepare for infection experiments, the 10403S strain of *L. monocytogenes* was streaked from a glycerol stock onto BHI plates supplemented with 200μg/mL streptomycin and 7.5μg/mL chloramphenicol. The plates were incubated at 37°C for 48 hours, after which bacteria were inoculated into 2mL BHI cultures containing the same antibiotics. Streaked plates were stored at 4°C and discarded after two weeks. Bacterial cultures were grown overnight in the dark at room temperature without agitation. When an OD_600_ of 0.8 was reached, the bacteria were washed twice with PBS and added to A431D cells as described in the infection assay section below.

Similar procedures were followed in the culturing of *L. monocytogenes* EGD-e, except that *L. monocytogenes* EGD-e::pAD_1_-cGFP was streaked onto BHI plates without antibiotic selection, while *L. monocytogenes* EGD-e *ΔagrD*::pAD_1_-cGFP was added to BHI plates that were supplemented with 10μg/mL chloramphenicol. Prior to infection, these two strains were cultured overnight in BHI media containing the same antibiotics.

### Mammalian Cell Culture

DMEM supplemented with high glucose (4.5g/L) (Thermofisher; 11965092), 10% fetal bovine serum (FBS) (GemBio; 900-108), antibiotic-antimycotic (Thermofisher; 15240096) and geneticin (800μg/mL) (Thermofisher; 10131035) was used to culture A431D human epidermoid carcinoma cells (Lewis et al., 1997) (gift from Cara Gottardi, Northwestern University).

### Retroviral Transduction of E-cadherin Constructs

As previously reported (Ortega et al., 2017), various E-cadherin constructs were designed using the full-length human E-cadherin sequence from the pcDNA3 human E-cadherin plasmid (a gift from Cara Gottardi’s lab at Northwestern University). Δcyto E-cadherin was generated by deleting amino acids 731-882 from the coding sequence for full-length E-cadherin, while the K738R, K816R E-cadherin construct was made by mutating two lysines at positions 738 and 816 to arginines. The WT, Δcyto and K738R, K816R E-cadherin constructs were cloned into pLNCX2 Retroviral Vector (Clontech) by Epoch Life Sciences Inc. GP2-293 cells (Clontech) were transfected with 10μg of one of the E-cadherin constructs as well as an amphotropic packaging vector (Clontech). The virus was concentrated 24 hours post transfection and filtered through a 0.45μm cellulose acetate filter 48 hours post transfection. It was then added to A431D cells along with 8μg/mL polybrene (Thermofisher; TR1003G) in the transduction step. Transduced A431D cells were selected for using media (high glucose DMEM, 10% FBS, antibiotic-antimycotic (ABAM)) supplemented with 800μg/mL of the selective antibiotic geneticin.

#### Infection Assay

Cleaned 18mm glass coverslips or 12-well glass-bottom plates (Cellvis; P12-1.5H-N) were pre-coated by incubation with 1µg/ml fibronectin overnight and then rinsed with PBS. A431D cells were seeded at a density of 5×10^5^ cells/mL and were allowed to incubate at 37°C for 36 hours. Experiments in which cells were fixed before image acquisition were conducted on glass coverslips (Supp. Fig. 1A and Fig. 4), while glass-bottom plates were used for live cell experiments (Fig. 1, Supp. Fig. 1B-G, Fig. 2, Supp. Fig. 2, Fig. 3, Supp. Fig. 3, Fig. 4, Supp. Fig. 4). Thirty-six hours after seeding, one mL of the bacterial culture at OD 0.8 was centrifuged for four minutes at 2000g. The BHI media was then aspirated and the pellet was washed twice with PBS before resuspension in DMEM. Host A431D cells were washed once with DMEM before the bacteria were added to them at a multiplicity of infection (MOI) of 200- 300 bacteria per host cell. Bacteria and host cells were incubated at 37°C for 10 minutes and then host cells were washed three times with DMEM to remove non-adherent bacteria. To perform live experiments, host cells were incubated in DMEM at 37°C for 15 minutes and subsequently incubated for an additional 15 hours in DMEM supplemented with 10% FBS and 20μg/mL gentamicin, an antibiotic which kills extracellular bacteria to prevent further invasion (Portnoy et al., 1988). The length of infection in the presence of gentamicin was 12 rather than 15 hours for experiments in which the samples were fixed. Drug treatments shown in Figure 4 included 5μM caveolin-scaffolding domain peptide (Sigma Aldrich Inc; 219482) 100ng/mL EGF (Sigma Aldrich Inc,. E9644) and10μM EIPA (Thermofisher; 337810). All drugs were added to infected host cells at the same time as gentamicin. Each fixed experiment was conducted on at least three experimental days using two coverslips per condition on each day.

#### Transfection of Plasmid DNA into A431D Cells

A431D cells were seeded on 6-well glass-bottom plates (Cellvis; P12-1.5H-N) that were pre-coated with 1μg/mL fibronectin at a density of 5×10^5^ cells/mL. The cells were allowed to incubate at 37°C for 24 hours after which full media was replaced with antibiotic-free media. In a separate 1.5mL microcentrifuge tube (USA Sci Inc; 1615-5510), FuGENE® Transfection Reagent (Promega; E2311) was added to Opti-MEM media (Thermofisher; 31985062) and subsequently mixed with 2μg each of pMD2.G, the VSV-G envelope expressing plasmid (Addgene; Plasmid #12259) and mEmerald-N1 (Addgene; Plasmid #53976). The transfection mix was incubated for 15 minutes at room temperature, as advised by the Fugene HD Transfection Reagent Technical Manual (Promega; Literature #E3211). Afterwards, the transfection mix was applied onto the monolayer of A431D cells and incubated for 12 hours at 37°C. Cytoplasmic fluorescence of mEmerald at 525nm upon illumination with 470nm light was used to identify transfected cells.

#### Time-Lapse Microscopy

WT, K738R, K816R and Δcyto E-cad A431D cells seeded onto fibronectin-coated glass bottom plates (Cellvis; P12-1.5H-N) were infected with *L. monocytogenes* as described above. At five hours post infection, infected cells were washed with L-15 (Thermofisher; 11415064) three times and host cell nuclei were then stained with Hoechst 33342 (Invitrogen; H3570) in L- 15 with 20μg/mL gentamicin for 15 minutes at 37°C. As imaging live cells in media containing phenol red introduces excessive background fluorescence to the acquired images, imaging media was prepared by supplementing DMEM lacking phenol red (Sigma Aldrich; D1145) with 4mM L-glutamine and 10% FBS. After three washes in L-15, cells were immersed in imaging media and images were collected every five minutes until 15 hours post infection using an inverted Nikon Eclipse TI2 with the perfect focus option selected, an EMCCD Camera (Andor Technologies) and a 20X 0.75 NA Plan Apo Air Objective. To ensure that only internalized bacteria were imaged, *L. monocytogenes* EGD-e::pAD_1_-cGFP and *L. monocytogenes* EGD-e *ΔagrD*::pAD_1_-cGFP, where cGFP was constitutively expressed, were washed nine times before imaging to remove surface-bound bacteria. They were also imaged at 18 rather than 15 hours post-infection, as the lag time before achieving maximum growth rate was longer for *L. monocytogenes* EGD-e as compared to 10403S. For the experiments in which A431D cells were infected with ActA mutant *L. monocytogenes*, images were acquired every 15 minutes to prevent phototoxicity, as transfected host cells exhibited additional sensitivity to fluorescent light. To observe infection of host cells in glucose-free media (Fig. 2A-B), a second imaging media was prepared using DMEM that did not contain glucose (Thermofisher; 11966025). All other components of the glucose-free imaging media were identical to the original and cells were immersed in it just prior to the start of the time-lapse recording at 5 h.p.i. Micromanager (Edelstein et al., 2014) was used to operate all microscopy equipment. An environmental chamber was used to maintain the temperature at 37°C and deliver 5% CO_2_ to the cells.

#### Fixation of A431D Cells and Labeling of Nuclei in Fixed Cells

For experiments in which the effect of a perturbation on efficiency of spread and bacterial load was assessed at a specific time point post-infection (Fig. 4), coverslips were fixed with 4% formaldehyde Thermofisher; 28906) for 10 minutes at room temperature, permeabilized with 0.1% Triton X-100 (Sigma Aldrich Inc; T9284) for an additional 10 minutes and stained with the DNA dye DAPI (Thermofisher; D1306), in blocking solution (2% BSA in PBS) for 45 minutes. Coverslips were then mounted onto slides using VECTASHIELD mounting medium (Vector Labs; H-1000) prior to imaging.

#### Partitioning of Each Bacterial Focus into Five Distinct Regions

We sought to divide each focus into a number of regions that could be compared across many foci collected over multiple independent experiments. Unlike focus size which varies widely, the length of the time lapse recording of bacterial focus growth can be kept constant.

Therefore, we partitioned the focus into distinct regions based on time of exposure to bacteria, where each compartment was bounded by the outlines of the bacterial focus set two hours apart. Fiji was used to crop each time-lapse recording of a bacterial focus such that extraneous bacteria were eliminated from the field of view. MATLAB_R2018B was then used to threshold and binarize the image of the focus. Using either the global threshold function graythresh or defining a specific threshold failed to account for the change in dynamic range of fluorescence intensity through the duration of the movie. At early time points, each focus had few dim bacteria that were equally bright but as the movie progressed, the brightness of the densely packed regions within a focus far exceeded the intensity of individual bacteria found at the edge. To be able to visualize peripheral bacteria in every frame, the MATLAB function multithresh was used to divide each focus into separate levels based on intensity and compute a specific threshold for each level. These multiple thresholds were then used to construct a binary image that more accurately represented bacterial positions than a binarization generated using a single threshold.

Binary images corresponding to the initial frame of the time-lapse recording to those depicting the focus at time points set two hours apart were summed to generate maximum intensity projections representing bacterial positions from 0 - 7 h.p.i., 0 - 9 h.p.i., 0 - 11 h.p.i. 0 - 13 h.p.i., and 0 - 15 h.p.i. We then generated an alphashape around each projection using the MATLAB function alphaShape with an alpha radius of 30 pixels and selecting the option to suppress all interior holes (Radhakrishnan et al., 2022) (Supp. Fig.2A). Each region was defined as the difference between two consecutive alphashapes and applied as a mask onto the greyscale image sequence (Supp. Fig.2A). Fluorescence intensity was quantified for every frame of the masked movie and used to calculate an inflection point for bacterial growth within each region.

#### Quantification of the Inflection Point at Which Bacterial Growth Deviates from Exponential Growth at a Constant Rate

Early in infection, bacterial replication can be modeled by exponential growth at a constant rate given by the equation 𝑁 = 𝑁_!_𝑒*^µ^*^"^, where 𝑁 represents the bacterial number at a given time, 𝑁_!_the initial number of bacteria per focus, *µ* the rate of growth expressed in minutes^-1^, and 𝑡 the number of minutes post infection. Similarly, when bacterial foci are partitioned as previously described and bacterial growth is monitored in the regime where the value of 𝑘 is dominated by growth rather than influx of bacteria into the region, bacterial behavior resembles exponential growth at a constant rate. In both instances, bacterial growth rate decreases over time. To determine the point at which bacterial growth rate began to slow in the focus as a whole, we fit multiple exponentials to the growth curve such that each exponential fit included an increasing number of data points (the first exponential included only fluorescence measured in the first frame of the time-lapse sequence, the second exponential incorporated fluorescence measured in the first two frames and so on). We extracted the growth rate constant, µ, from each of these exponential fits and found the time after which its value decreased monotonically. We called this time the inflection point T_i_ and used the corresponding fit to describe the portion of the growth curve exhibiting exponential growth at a constant rate. To determine the inflection point for bacterial growth rate in separate regions within a bacterial focus, all of the data points associated with times in which there were either no bacteria within the region or growth rate was dominated by the bacterial invasion into the region, were excluded from the analysis. Then, a similar procedure involving calculating the time after which *µ* consistently decreased was followed to determine the inflection point at which the slowing of bacterial growth rate commenced. We did not report an inflection point for the outermost region in between focus boundaries drawn at 13 and 15 h.p.i., as the growth rate of bacteria populating this region did not begin to slow by the end of the time-lapse movie at 15 h.p.i. for the vast majority of foci analyzed. (Fig. 1I).

#### Quantification of Bacterial Growth Rate over Time

Bacterial growth rate over time was quantified by fitting distinct exponential functions to the growth curve at 1 hour time intervals (Supp. Fig. 1C). The value of *µ* for each exponential was designated as the growth rate within the time interval. Doubling time was determined by dividing the growth rate from the natural log of 2.

#### Determination of Time Required for µ to Accurately Reflect Growth Rate in Each Region of a Bacterial Focus

As described previously, each focus was partitioned into five regions. Bacterial growth rate upon entry into every region other than the center was characterized by an initial spike during which the value of *µ* exceeded the expected 0.009 min^-1^ followed by exponential growth at a constant rate and then an ultimate slowing of bacterial growth rate.

We ran a stochastic simulation of *L. monocytogenes* cell-to-cell spread (Ortega et al., 2019) to determine whether the unusually high value of *µ* upon entry into a new region depended on an increase in the rate of bacterial replication. To do so, we first adapted the input parameters of diffusivity and replication rate in the simulation to closely reflect experimental values. In particular, the rate of bacterial replication was kept at a constant 0.01 minutes^-1^ throughout 15 hours of infection. We then partitioned each focus into five regions as described previously and used the MATLAB function inShape to count the number of bacteria within each region at every timestep of the simulation. When we plotted bacterial number (Supp. Fig. 2B) and growth rate (Supp. Fig. 2C) over time, we found that *µ* remained constant in the most interior region but was initially much higher than the input value for every other region of the simulated focus (Supp. Fig. 2B-C). This result suggested that the early spike in growth rate was a consequence of bacterial flux into a new compartment of the focus rather than an outcome caused by a genuine increase in rate of bacterial replication.

Using the simulation for bacterial spread, we then determined the time after entry into each exterior region at which bacterial behavior returned to exponential growth at a constant rate (Supp. Fig. 2C). This time was defined as the first timestep at which the bacterial number was equal to or higher than the corresponding number for the exponential fits to the growth curves (depicted as the grey dashed lines in Supp. Fig. 2E) for 10 consecutive timesteps. We then discarded experimental data points (such as that shown in Fig. 1J) that fell before these determined times and fit exponentials to the remaining growth curves to find the inflection point for bacterial growth in each region as previously described (Fig. 1K, Fig. 3I, Supp. Fig. 4C).

#### Image Acquisition of Fixed Samples Using Confocal Microscopy

We acquired confocal images of mTagRFP expressing *L. monocytogenes* as well as DAPI-stained host cell nuclei. A z-spacing of 1μm was used to image these foci and approximately 10 slices per infection focus were collected. These images were taken using a Yokogawa W1 Spinning Disk Confocal with Borealis Upgrade on a Nikon Eclipse Ti2 inverted microscope with a 50μm disk pattern (Andor). Additionally, a Plan Apo 20X 0.95 NA water immersion objective was used to acquire the images, a piezo z-stage (Ludl 96A600) was used to increase the rate of image acquisition between z-frames and MicroManager v.1.4.23 (Edelstein et al., 2014) was used to operate all microscopy equipment. For the focus size experiments shown in Figure 4, up to fifty bacterial foci were imaged on each coverslip with two replicate coverslips per condition and each experiment repeated on three separate days.

#### Quantification of Bacterial Focus Area

After performing a maximum intensity projection of the confocal images used to generate data shown in Figure 4, we cropped the bacterial focus to eliminate extraneous bacteria in Fiji. Then, MATLAB_R2018B was used to threshold and binarize the image of the focus.

Using the MATLAB function alphaShape and choosing the option to fill interior holes, we generated an alpha shape with an alpha radius of 30 around each binarized focus and calculated its area. (Radhakrishnan et al., 2022)

A similar protocol was used to calculate the area of bacterial foci at 15 h.p.i. from the time-lapse recordings of bacterial growth (Fig. 3C), except that the analysis was performed exclusively on the last frame of the time-lapse movie and it was not necessary to calculate a maximum intensity projection since bacteria were imaged at a single focal plane using wide- field microscopy.

To calculate host cell area at one hour time points through a time-lapse recording (Fig. 4D-F), we binarized the mEmerald-N1 signal which filled the cytoplasm of the host cell. We then generated an alpha shape around the binarized image, using an alpha value of 5 rather than 30 to account for the differences in cell size and shape between a bacterial focus and single mammalian cell. Calculation of cell area was as previously described.

#### Quantification of Bacterial Fluorescence Intensity

Bacterial fluorescence intensity was quantified as previously described (Ortega et al., 2019). Briefly, the focus was binarized at each time point and the mask of the resulting image was dilated until the focus was depicted as one continuous shape. Then, the median intensity of all points in the field of view that were excluded from the dilated mask was calculated and determined to be the background fluorescence intensity. This value was subtracted from all pixel values in the original greyscale image. Intensity values for all pixels in the now background-subtracted image were averaged to produce the fluorescence intensity of the bacterial focus.

As host cells were incubated with ActA mutant *L. monocytogenes* for a longer duration before gentamicin addition as compared to the WT bacterium, many more host cells were infected with bacteria. I applied a mask of the host cell area (obtained by generating an alphashape around the cytoplasmic mEmerald-N1 signal) onto the RFP channel containing bacteria. In this way bacterial fluorescence from neighboring host cells was discarded, allowing for the quantification of fluorescence from a single host cell of interest. Other steps in the calculation of fluorescence intensity were as listed above, except that intensity values of all pixels were added rather than averaged to allow for the calculation of total number of bacteria per cell and the carrying capacity of the cytoplasm.

Fitting Growth Curve of actA mutant 34 L. monocytogenes to the Gompertz Function:

Intracellular growth of mTagRFP actA mutant 34 *L. monocytogenes*, a bacterial mutant which is unable to spread from cell to cell, was monitored from 5 h.p.i. to 15 h.p.i. through microscopy. A growth curve was obtained by fitting the trajectory of bacterial growth over time to the form of the Gompertz function given below.

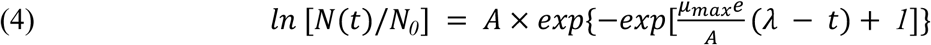

Due to the difficulty of distinguishing individual bacteria within the host cell cytoplasm, the fluorescence intensity of the bacterium was used as a substitute for bacterial number (𝑁).

Furthermore, since the mammalian host cell grows over time, the Gompertz function was fit to the plot of density (fluorescence intensity divided by the area of the mammalian host cell) against time rather than raw fluorescence intensity value itself. The maximal growth rate (𝜇_’()_) was calculated as previously described, by fitting an exponential function to consecutive data points from the beginning of the time course, at 5 h.p.i., to 𝑇_&_ or the inflection point after which growth rate begins to decrease monotonically. The value of 𝜆 or lag time was then solved for using the following equation.

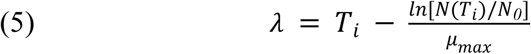

The remaining constant 𝐴, defined as the carrying capacity, was determined by performing a fit to the experimental data using the MATLAB package, cftool.

#### Quantification and Statistical Analysis

The number of data points evaluated for each experiment as well as statistical parameters and significance are reported in the Figures and the Figure Legends.

Times of inflection for various regions within a bacterial focus (Fig. 1K, Fig. 3I and Supp. Fig. 4C) were represented as bar graphs with the midlines signifying the mean of the data points and the whiskers denoting the standard deviation. Fluorescence intensity over time (Fig. 1E-F) and growth rate over time (Fig. 1G, Fig. 2B-C, Fig. 3E, Supp. Fig. 3, Supp. Fig. 4A) were depicted as line plots with midlines representing the mean of the datapoints for each condition, error bars representing the standard error of the mean and the shaded regions above and below indicating the standard deviation. Fluorescence intensity over time for representative foci, simulated foci or distinct regions within a bacterial focus (Figures 1I, Supp. Fig. 1B, Supp. Fig. 1E-G) and growth rate over time for distinct regions within a bacterial focus (Figures 1J, Supp. Fig. 2C) were depicted in a similar manner, except that standard deviations were not included in the figure. P-values were calculated using the Wilcoxon rank-sum test in GraphPad PRISM8 with a threshold of significance set at 0.05. For plots of growth rate as a function of time, asterisks or the abbreviation “ns” were placed above each datapoint to denote the significance of the difference in growth rate between the two conditions within a shared time interval. The table below translates the meaning of each asterisk.

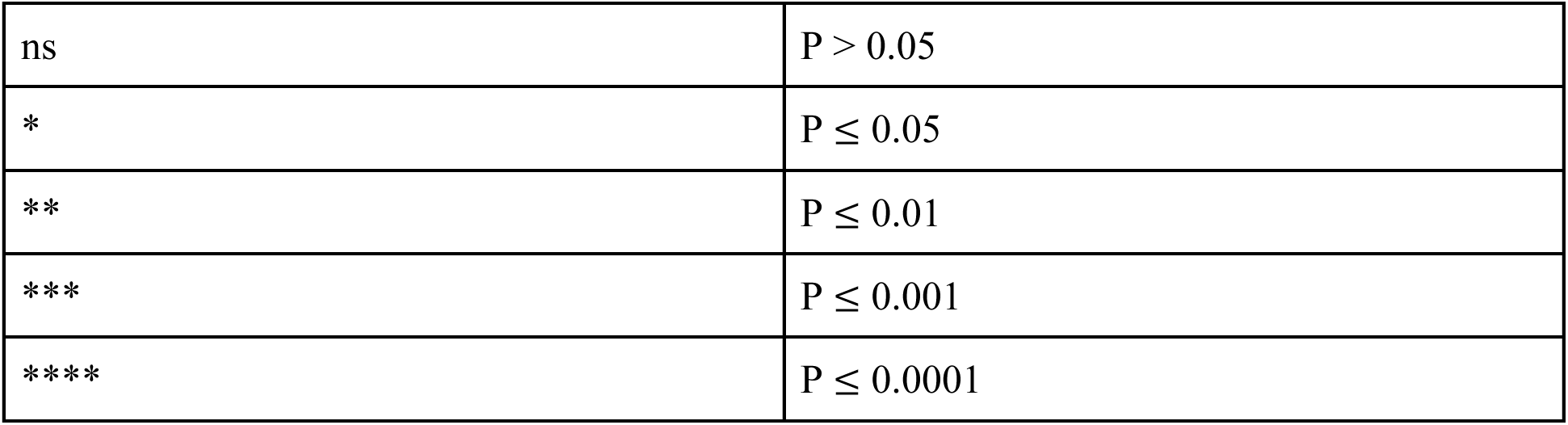

|  |  |
| --- | --- |
| ns | $P > 0.05$ |
| * | $P \leq 0.05$ |
| ** | $P \leq 0.01$ |
| *** | $P \leq 0.001$ |
| **** | $P \leq 0.0001$ |

Time of inflection for the focus as a whole in Fig. 1H, Fig. 3F and Supp. Fig. 4C, normalized focus area in Fig. 3C and Fig. 4A and normalized fluorescence intensity in Fig. 3D and Supp. Fig. 4B were represented as violin plots. In the violin plots, the dashed line denotes the median of the distribution and the dotted lines represent the 25 and 75% quartile of the pooled data from all independent experiments. The mean for each individual experiment is indicated within the violins as a filled circle. P-values for the comparison of cell-to-cell spread efficiency were calculated using the linear mixed-effects model (Gałecki and Burzykowski, 2013), which takes into account the variability introduced by random effects such as the day of experimentation and coverslip on which cells were seeded. As at least three independent experiments were conducted to evaluate whether a perturbation affected focus area and fluorescence intensity, the threshold for significance was set at 0.01 to account for multiple hypothesis testing.

---

**Supplemental Figure 1.**
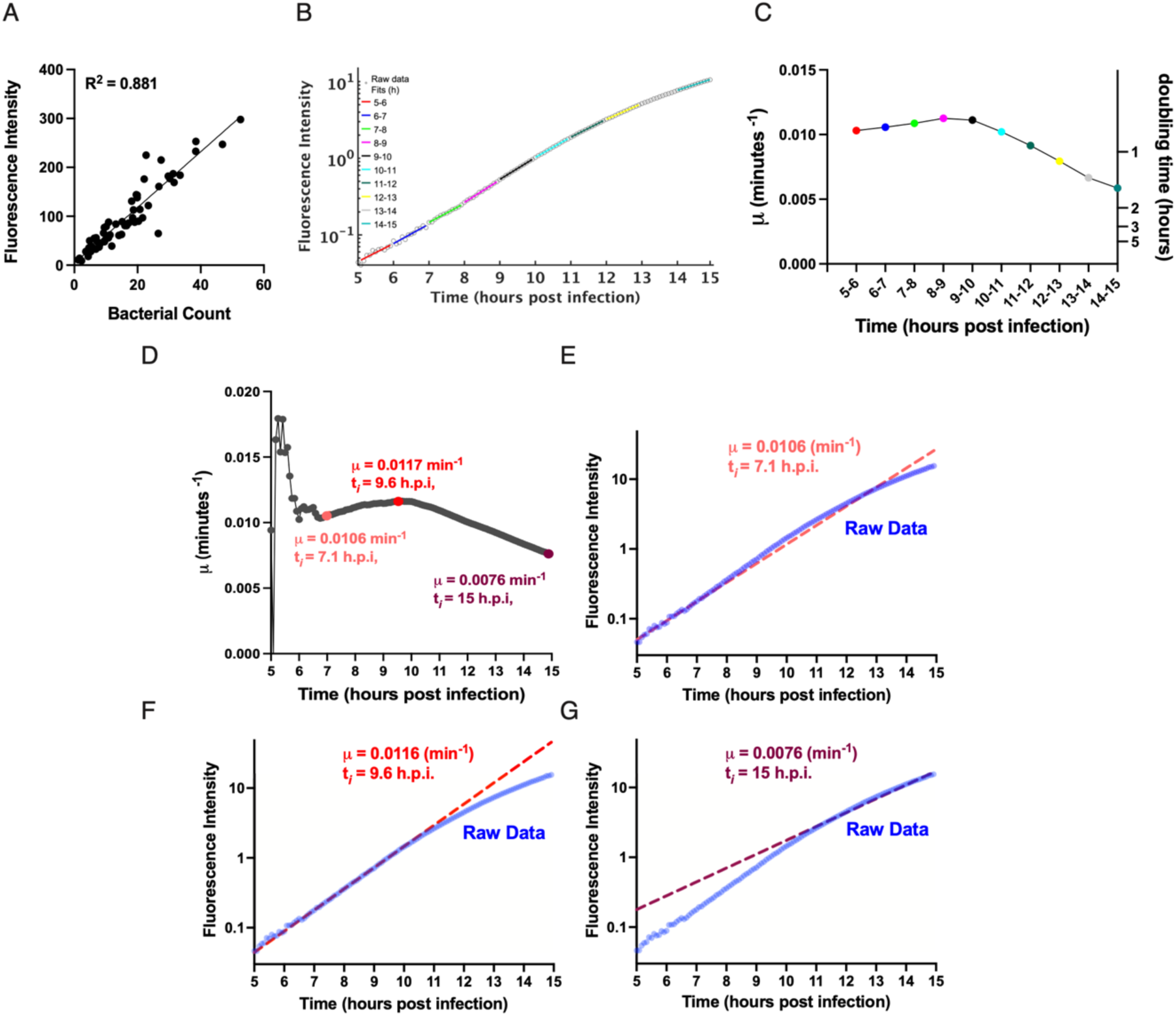
Growth rate µ and inflection time T_i_ for intracellular replication of *L. monocytogenes* can be determined directly from measurements of bacterial fluorescence intensity over time. **A.** Correlation between number of bacteria in a focus and fluorescence intensity at 6 hours post infection. 62 foci were imaged for this analysis. **B.** Fluorescence intensity plotted as a function of time for a representative bacterial focus imaged from 5 h.p.i. to 15 h.p.i. Open circles represent individual fluorescence intensity measurements, while colored lines depict the exponential fit to the underlying curve at one hour time intervals. **C.** Growth rate plotted as a function of time at one hour time intervals for the bacterial growth curve shown in B. Each data point represents the growth rate calculated from the corresponding exponential fit in B, where the color of each data point in C. matches the color of its respective fit line in B. **D.** Growth rate µ calculated by fitting an exponential function to only the subset of a representative bacterial growth curve from the beginning of the measurement, where integrated time or t_i_ = 5 h.p.i., to the time corresponding to the value of the x-axis at each data point. Three specific points on the curve are highlighted in salmon, red and maroon and depict growth constants obtained using fits from t_i_ = 5 h.p.i. to t_i_ = 7.1 h.p.i., t_i_ = 9.6 h.p.i. And t_i_ = 15 h.p.i. respectively. E. - G. Remaining graphs show the plot of fluorescence intensity over time for the length of the experiment (blue curve) against an exponential fit to all data points collected from (E) 5 h.p.i. to 7.1 h.p.i. (salmon dashed line), (F) 9.6 h.p.i. (red dashed line) or (G) 15 h.p.i. (maroon dashed line).

**Supplemental Figure 2:**
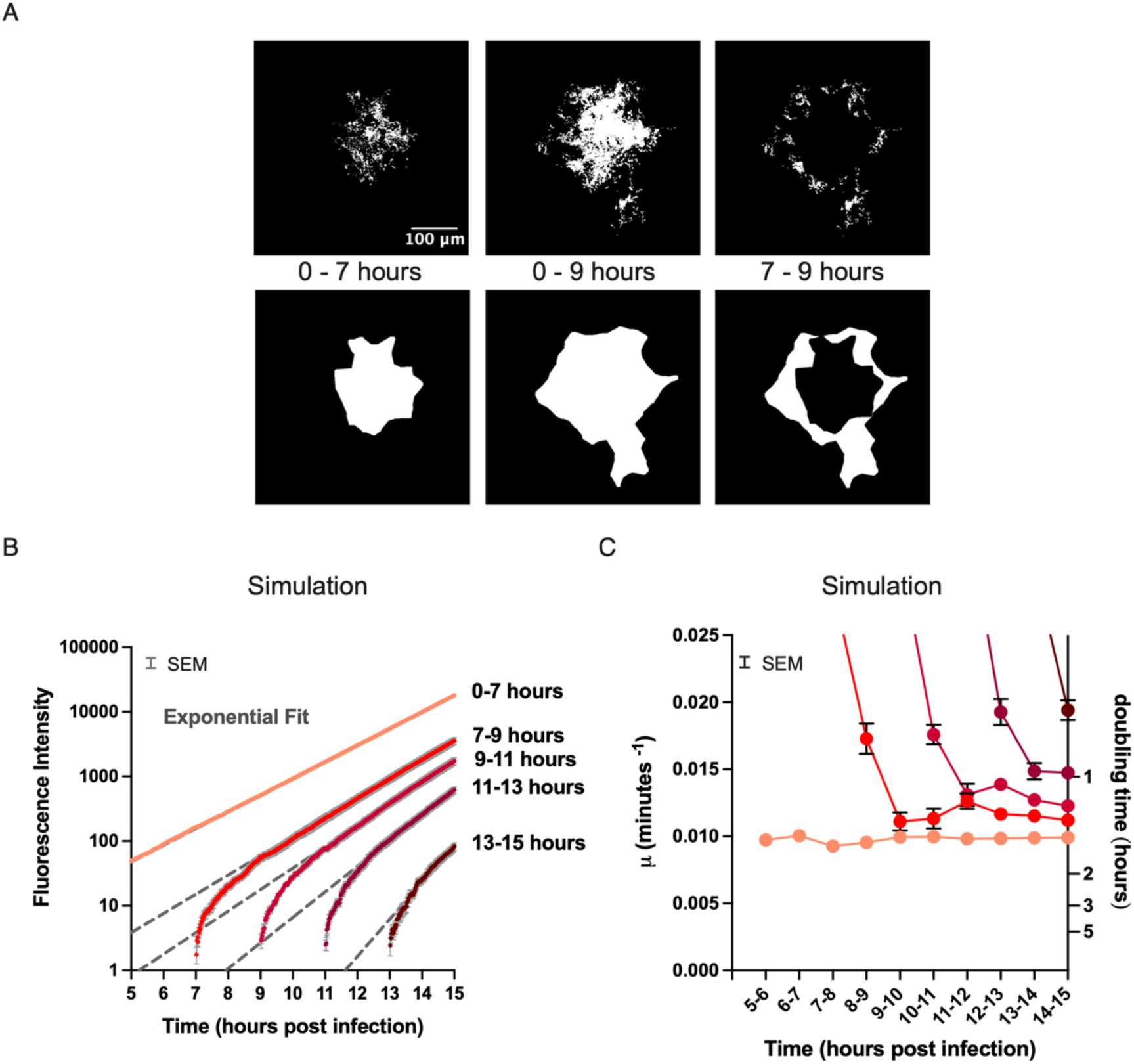
Stochastic simulations of *L. monocytogenes* cell-to-cell spread aid in determining inflection point for distinct regions of a bacterial focus. **A.** To partition a focus into 5 regions, alpha shapes were first generated to bound the maximum intensity projection of binarized bacteria from the beginning of the time lapse recording to time points set 2 hours apart, such as 7 and 9 hours post infection. The difference between two consecutive projections yielded a region within which bacterial growth could be quantified. **B**. Stochastic simulations of bacterial spread (Ortega et al., 2021) were used to generate realistic simulated foci matching the average growth characteristics of the experimental data, and were analyzed using the same methods. Bacterial replication was set to occur at a constant rate for the duration of each simulation. Quantification of average bacterial fluorescence intensity is plotted as a function of time within distinct regions of a simulated focus. Dashed lines depict the exponential fits to each growth curve. **C.** Growth rate µ of simulated bacteria as a function of time within distinct regions of a simulated bacterial focus, averaged over 10 simulated foci. For B. and C., the color of the curves indicates the region within which bacterial growth is monitored, and the error bars depict the standard error of the mean.

**Supplemental Figure 3:**
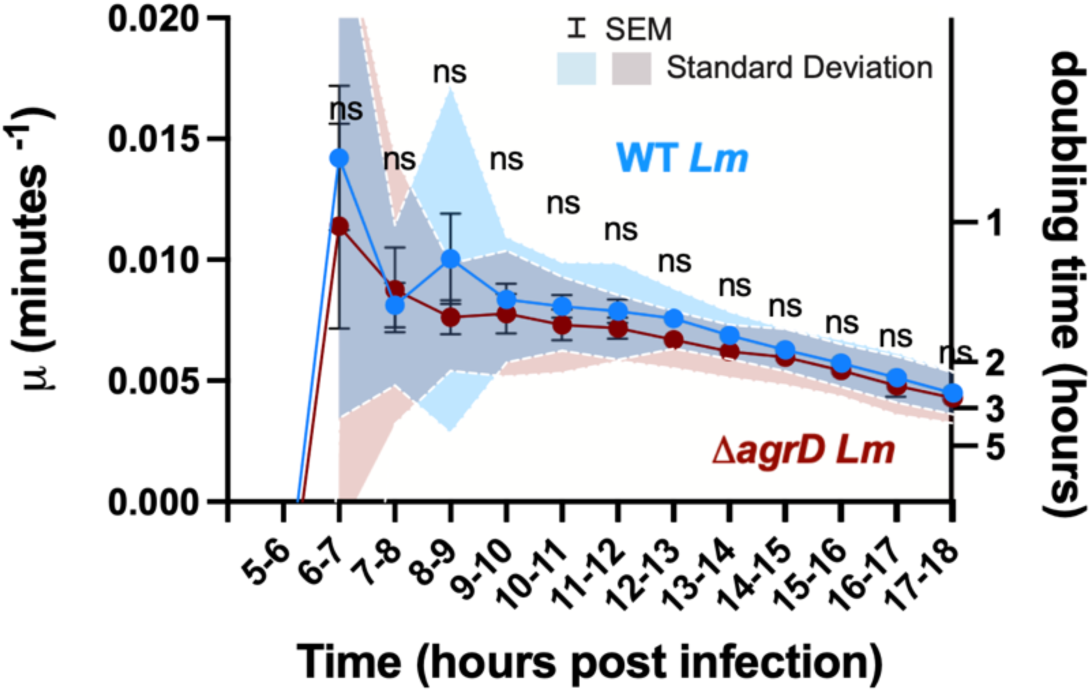
Quorum sensing among bacteria does not contribute to the decrease in growth rate of *Listeria monocytogenes* over time. Growth rate as a function of time for cells WT *L. monocytogenes* EGD-e (light blue trace) and *ΔagrD L. monocytogenes* EGD-e (brown trace). The number of foci imaged for each condition include 17 and 10 foci respectively. P values were calculated using the Wilcoxon Rank-Sum test and denoted as asterisks above each data point. The error bars represent the standard error of the mean and the shaded region the standard deviation.

**Supplemental Figure 4:**
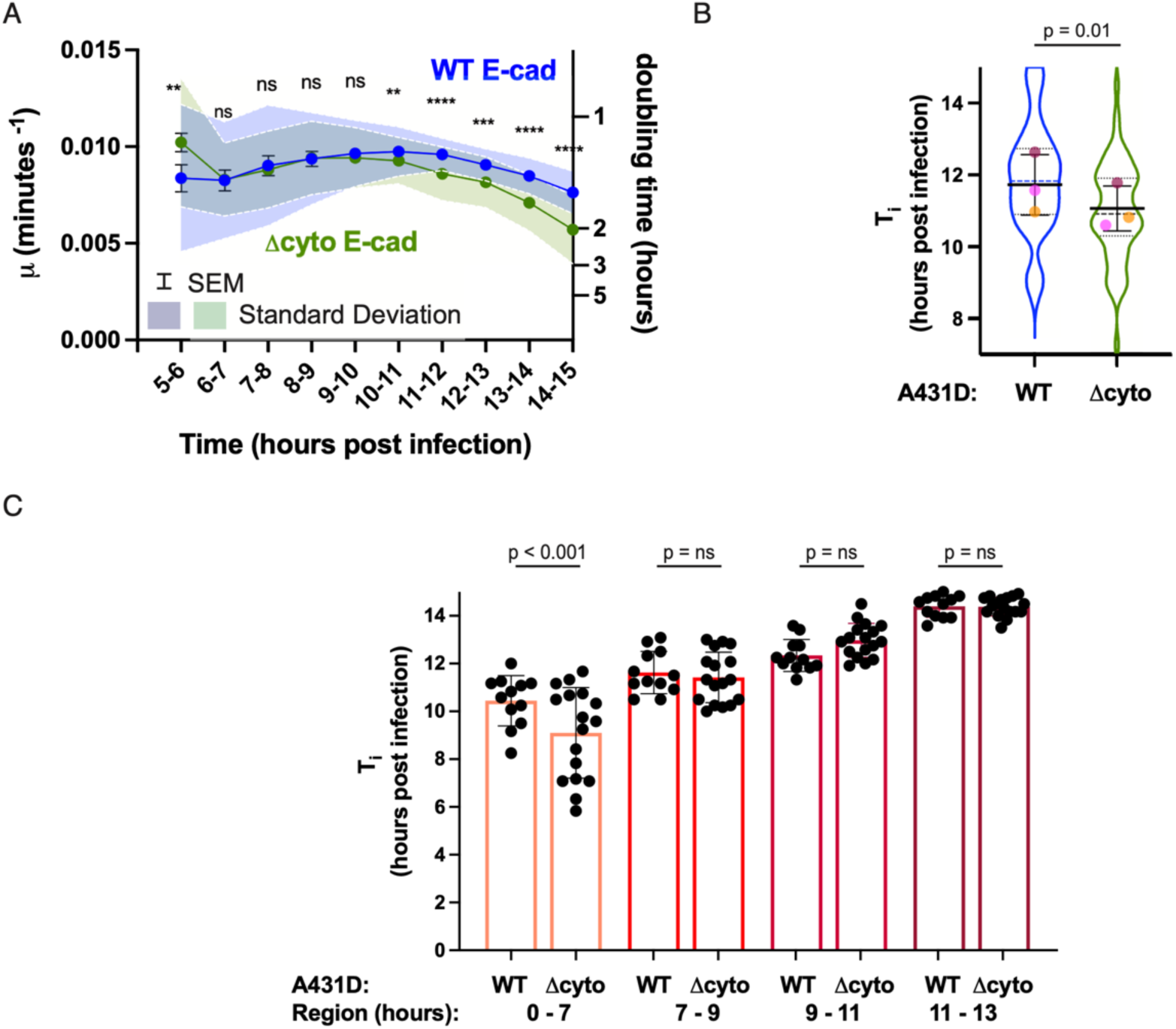
Removal of the cytoplasmic domain of host cell E-cadherin reduces *L. monocytogenes* cell-to-cell spread and causes the growth rate inflection point to occur at an earlier time. **A.** Growth rate µ as a function of time for bacteria propagating through WT E-cad (blue trace) and Δcyto E-cad (green trace) A431D cells. P values were calculated using the Wilcoxon Rank-Sum test and denoted as asterisks above each data point. Error bars represent the standard error of the mean and shaded region the standard deviation. 49 and 45 foci were imaged for each condition respectively. **B.** Comparison between inflection point T_i_ for bacteria replicating in WT E-cad or and Δcyto E-cad cells. P values were determined using the linear mixed-effects model. **C**. Inflection point at which bacterial growth begins to slow in each region of the focus in WT or Δcyto E-cad A431D cells. The color of each bar corresponds to the region on the schematics in Figure 1D. P-values were calculated using the Wilcoxon rank-sum test. 15 foci were included in the analysis for each condition.

